# Independent transposon exaptation is a widespread mechanism of shadow enhancer evolution in the mammalian genome

**DOI:** 10.1101/703314

**Authors:** Nicolai K. H. Barth, Lifei Li, Leila Taher

## Abstract

Many regulatory networks involve (possibly partially) redundant *cis*-regulatory elements. These elements are commonly referred to as shadow enhancers and have been assumed to mainly originate by sequence duplication. An alternative mechanism that has not been examined involves the independent exaptation of transposons. Here, we identified 2,117 and 2,787 pairs of shadow enhancers in the human and mouse genomes, respectively, and found that 34% and 18% of these pairs derive from transposons. Moreover, we show that virtually all the enhancer partners of these transposon-derived shadow enhancer pairs are independent exaptations from different transposon types and hypothesize that the increase in the expression and tissue-specificity of their targets as compared to those of non-shadow enhancers facilitates their fixation in the genome. Finally, we provide evidence that the regulatory redundancy observed in many mammalian regulatory networks is predominantly driven by convergent evolution, since shadow enhancers are hardly ever conserved across species.

## Introduction

Already before the DNA-sequencing era, changes in gene regulation have been proposed as the most likely explanation for most phenotypic discrepancies between both individuals of the same species and closely related species [1]. In Eukaryotes, gene expression is primarily regulated at the level of transcription. Transcription is initiated when the RNA polymerase II machinery recognizes and binds specific sequences in the core promoter of a gene. The resulting basal level of expression can be increased or decreased through biochemical interactions between transcription factor (TF) and co-factor proteins and *cis*-regulatory elements scattered throughout the genome. *Cis*-regulatory elements include sequences that are proximal to their target genes, such as promoters, and sequences that are distal, such as enhancers. These elements act together to control the expression of their target gene(s). Current data suggest that most genetic variants associated with complex disease are located within regulatory elements as opposed to genes [2,3].

Enhancers are typically a few hundred base pairs long and harbor clusters of transcription factor binding sites (TFBSs). They are usually bound by tissue-specific TFs and can thereby produce highly controlled spatiotemporal gene expression patterns [4]. The expression of many genes has been shown to be controlled by two or more – rather than one – enhancers with overlapping regulatory activities [5–11]. Such regulatory redundancy has been suggested as a means to impart robustness and ensure steady gene expression, for example, under adverse environmental or genetic conditions [12–14]. In 2008 Mike Levine and colleagues coined the expression “shadow enhancers” to refer to enhancers with redundant regulatory activities in the fruit fly *Drosophila melanogaster* [5]. Specifically, they found that the genes *vnd* and *miR-1* are both regulated by at least a pair of enhancers each. One of the enhancers in these pairs – the “primary” enhancer – is more proximal to the TSS of their target than the other – the “shadow” enhancer – but both enhancers have similar regulatory activities and bind the same TFs. Since then, the expression “shadow enhancers” has been extended to a more general use to describe two or more (possibly partially) redundant enhancers, bypassing the assignment of the labels “primary” and “shadow”. Here, we apply the expression “shadow enhancers” to two or more enhancers with (possibly partially) redundant regulatory activities, i.e. controlling the same gene(s) in the same tissue(s). In that sense, recent work has suggested that shadow enhancers are common in the mammalian genome [13].

The acquisition of new enhancers is thought to play a key role in gene neo-functionalization and to be a major evolutionary force contributing to the remarkable diversity of mammals and other species [15]. Shadow enhancers in the human genome may have originated through several mechanisms, including sequence duplication and independent exaptation or co-option of transposons [15]. In particular, much evidence has accumulated suggesting exaptation of transposons as enhancers [16]. Transposons are mobile genetic sequences that can jump around the genome from one location to another, behaving as genomic parasites. They have been very effective in colonizing many mammalian and non-mammalian genomes, and occupy nearly half of the human genome [17,18]. Transposons harbor TFBSs and their insertion has been shown to influence the expression of nearby genes in reporter gene assays [19–25] and also using the CRISPRcas9 system [26]. Moreover, it has been shown that groups of transposons located close to each other may evolve to (synergistically) work together to ensure robust gene expression within correct spatiotemporal limits [23]. In addition, Franchini et al showed that two enhancers that redundantly control the expression of the proopiomelanocortin gene (POMC) in a specific group of neurons originated from the exaptation of two different retrotransposons: an LTR transposon between the metatherian/eutherian split (147 Mya) and the placental mammal radiation (~90 Mya), and a SINE retrotransposon before the origin of prototherians (166 Mya)[27]. The fact that two enhancers with overlapping regulatory activities are under purifying selection would be explained by their role as regulatory buffer that prevent deleterious phenotypic consequences upon the loss of one of them [13]. This is in agreement with the theory independently proposed by Schmalhausen and Waddington, which states that phenotypes will remain relatively invariant to perturbations [28].

In contrast to promoters, which are directly upstream of their target genes, enhancers can be located anywhere in the genome – upstream or downstream of genes, but also within introns and not necessarily of their target genes, making their identification and characterization challenging. Nevertheless, much progress has been made towards creating a catalogue of the regulatory elements in the human genome, in particular through chromatin immunoprecipitation (ChIP)-based methods and international consortia such as the Encyclopedia of DNA Elements (ENCODE, [29]). These data have made evident the pervasiveness of multiple enhancers with similar activities near the same gene [18] and are starting to reveal their adaptive value [13]. Nevertheless, it remains unknown how shadow enhancers originate. To directly assess this, we used cap analysis of gene expression (CAGE) data from the FANTOM project [30,31] and a stringent approach to identify 1,280 shadow enhancers in the human genome and 1,939 in the mouse genome, associated with 616 and 1,199 genes, respectively. By combining the transposon annotation of these shadow enhancers with phylogenetic information we provide evidence that in the vast majority of the shadow enhancer pairs where both partners are likely to result from transposon exaptations – 34% (18% in mouse) of the total – the enhancer partners have been acquired independently, throughout many periods of mammalian evolution, indicating that independent transposon exaptation is a widespread mechanism of shadow enhancer origination. Moreover, we found that shadow enhancers are highly lineage-specific, in the sense they are not evolutionary conserved, and that the similar levels of redundancy observed between orthologous mammalian regulatory networks are often examples of convergent evolution.

## Results

### Shadow enhancers are a common feature of human regulatory networks and have distinct features

Examples of enhancers with (possibly partially) redundant regulatory activities– commonly known as “shadow enhancers” – have been known for many years [32]. However, the extent to which such regulatory redundancy contributes to the robustness of mammalian regulatory networks is only now beginning to be appreciated [13]. To quantify partial and absolute regulatory redundancy in the human genome we analyzed a dataset of 201,799 and 54,284 putative promoters and enhancers, respectively, identified by the FANTOM5 Consortium with the CAGE technique ([30,31], see Methods). Active enhancers and promoters are often transcribed [33], and the levels and directionality of their transcription have been shown to reflect their activities [31]. Moreover, co-transcription of enhancers and promoters has been proposed as a means to associate enhancers with their target genes [31,34]. Finally, enhancers and their target genes are generally located within the same topologically associating domain (TAD), as TADs are organizational chromatin units with frequent internal interactions [35]. Thus, to uncover the enhancer target genes, we computed the Pearson correlation coefficient (*r*) between the transcriptional activity profiles of pairs of enhancers and promoters located within the same TADs across 44 groups of samples from related tissues – to which we refer as “facets” (see Methods) [31]. Of the enhancers predicted by the FANTOM5 consortium, 11,582 were active in at least one of the facets included in this study; in turn, 10,445 of these enhancers were located within a TAD that also comprised one or more active promoters. Of the promoters, 72,272 out of a total of 201,799 were active in at least one facet and located within 500 bp to a TSS of an ENSEMBL protein coding gene; 55,612 were located within TADs with at least one active enhancer. This yielded a total of 314,746 possible pair-wise associations between enhancers and promoters in 2,449 TADs. Nearly 3.5% of those pairs (10,952), involving 3,523 enhancers and 6,474 promoters in 1,398 TADs, were positively correlated (r > 0.5, P-value ≤ 0.0025, adjusted P-value < 0.05). We regarded the transcripts (genes) corresponding to the promoters (see Methods and Figure 1A) as the putative target transcripts (genes) of the enhancers. Based on this, we identified 2,117 pairs of enhancers with the exact same target transcripts and common activity in at least one facet. We further refer to these putative pairs of enhancers with (possibly partially) redundant regulatory activities as “shadow enhancer pairs” and to the partners of the pairs as “shadow enhancers”. Approximately 36% (1,280) of the enhancers that were correlated with at least one promoter were shadow enhancers. This result points towards enhancer redundancy being a widespread feature of regulatory networks and is in accordance with previous observations in the Drosophila genome [18].

**Figure 1.**
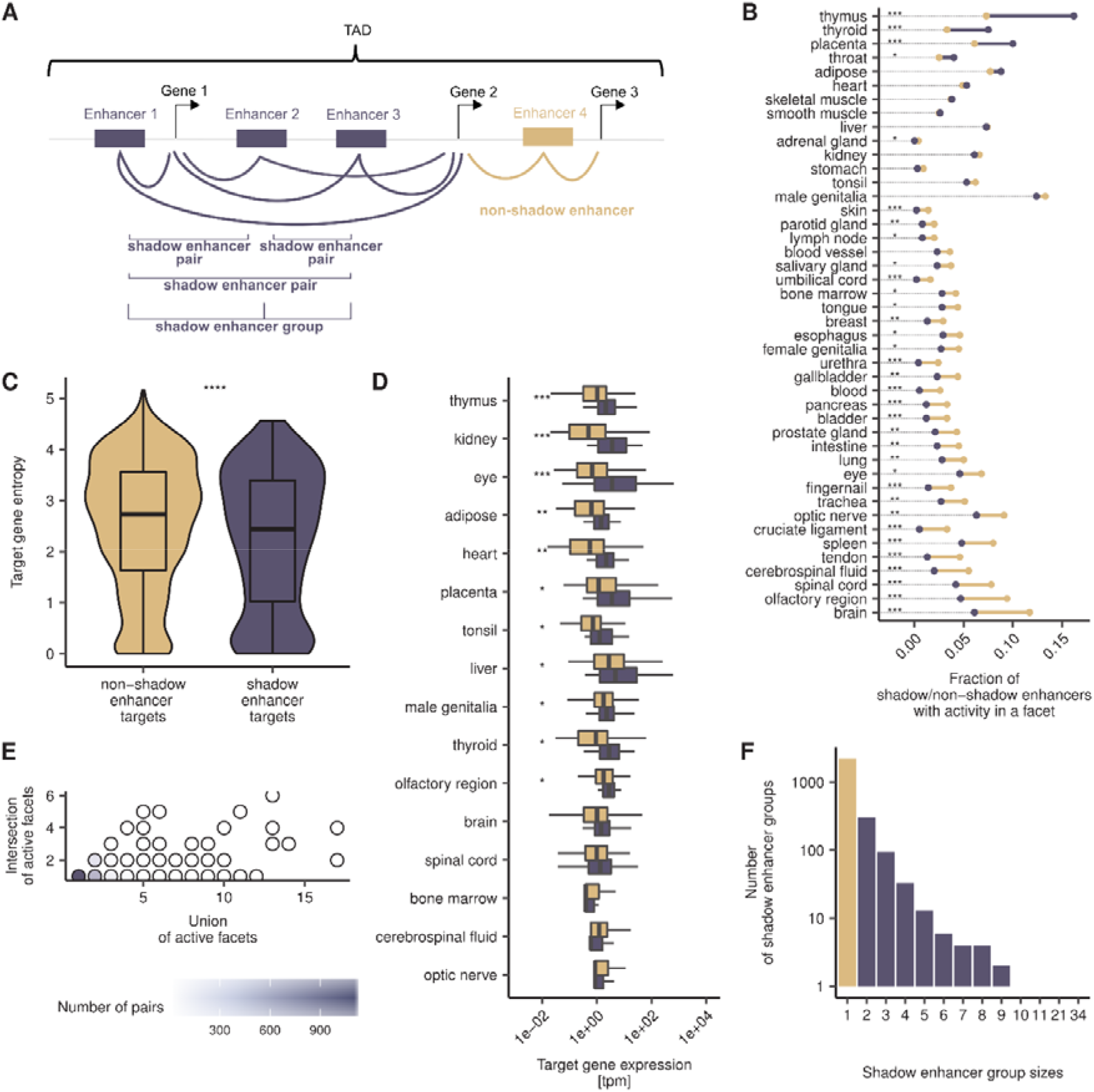
Shadow enhancers show distinctive features. **A)** Enhancers are assigned to target genes, further categorized into shadow and non-shadow enhancers, arranged into shadow enhancer pairs and shadow enhancer groups. If the activity profiles of two enhancers within a TAD are moderately significantly correlated with the activity profiles of the same set of promoters, then the two enhancers are considered a “shadow enhancer pair” and the genes associated with the promoters, their target genes. Significant correlations are shown as lines between enhancers and genes. All shadow enhancer pairs associated with the same target genes form a shadow enhancer group. **B)** Some facets show strong enrichments for active shadow enhancers (e.g., thymus), some show strong depletions (e.g., brain). The dumbbell plot shows the fractions of active non-shadow (beige) and shadow (purple) enhancers per facet. The facets are sorted according to the difference in the fractions of shadow and non-shadow enhancers; if the fraction of active non-shadow enhancers is larger than that of shadow enhancers, the line between the dots is depicted in beige; otherwise, in purple. The dotted lines serve as visual aids. Asterisks indicate adjusted P-values of Fisher’s exact tests: *’ P < 0.05, ’**’ P < 0.01, ’***’ P < 0.001 **C)** Shadow enhancer target genes are more tissue-specific than non-shadow enhancer target genes. Depicted are the entropies of non-shadow (beige) and shadow (purple) enhancer target genes. Asterisks indicate P-value of Wilcoxon rank sum test. ’****’ P < 0.0001. **D)** In most facets, shadow enhancer target genes show a stronger expression than non-shadow enhancer target genes. Facet-specific expression of non-shadow (beige) and shadow (purple) enhancer target genes with adjusted P-values of Wilcoxon rank sum tests. The facets are sorted according to adjusted P-values. See B) for significance code. **E)** The majority of shadow enhancer pairs have a 100% overlapping activity pattern. Regulatory redundancy of shadow enhancer pairs, measured as the ratio of the number of facets with shared activity to the number of facets in which any of the partners of the pair is active. **F)** Shadow enhancers form groups of different sizes. Depicted are the number of non-shadow enhancers (beige) and shadow (purple) enhancer in a group.

Both shadow and non-shadow enhancers were relatively weakly conserved (medians of average base-wise PhyloP scores of 0.04, see Methods). However, shadow enhancers had a higher SNP density than non-shadow enhancers (medians 110 SNPs/kb and 108 SNPs/kb, respectively, P-value = 0.01, two-sided Wilcoxon rank-sum test), suggesting differences in selective pressure within the human population. In addition, shadow enhancers and non-shadow enhancers had a similar number of target genes (medians = 1) and similar GC-content (medians = 0.48). We observed, however, that the distance of shadow enhancers to the nearest TSS of a target gene was smaller than for non-shadow enhancers (median distances 73 kb and 90 kb, respectively, P-value = 0.02, two-sided Wilcoxon rank-sum test, see Figure S1) and the target locus size (median value of all target loci of an enhancer) was larger (136 kb and 195 kb, P-value = 1.3 × 10^−12^two-sided Wilcoxon rank sum test). These results point towards shadow enhancers being preferentially located in less gene-dense regions. Shadow and non-shadow enhancers exhibited differences in their activities (see Figure 1B). Thus, compared to non-shadow enhancers, shadow enhancers where over-represented in facets such as thymus (adjusted P-value = 2.4×10^−14^, see Methods) and under-represented in brain (adjusted P-value = 4.3×10^−7^), cruciate ligament (adjusted P-value = 2.9×10^−7^), cerebrospinal fluid (adjusted P-value = 8.7×10^−7^) and olfactory region (adjusted P-value = 1.5×10^−6^), among other facets, suggesting that shadow enhancers are particularly important for certain tissues. Moreover, the target genes of shadow enhancers were more tissue-specific than those of non-shadow enhancers (median entropies 2.4 and 2.7, respectively; P-value = 4.2 × 10^−5^, two-sided Wilcoxon rank-sum test, see Figure 1C and Methods). Indeed, shadow enhancers are hypothesized to be associated with tissue specific expression patterns, whereas non-shadow enhancers are associated with unspecifically expressed house-keeping genes [13]. Finally, in agreement with findings in other tissues [36], we found that the expression strength of shadow enhancer target genes was generally stronger than that of non-shadow enhancer target genes (see Figure 1D). Specifically, it was stronger in eleven out of sixteen facets with enough enhancers for testing (adjusted P-values ≤ 0.05) and as strong in the remaining five facets (see Methods). Hence, enhancer redundancy could be a mechanism to increase gene expression levels – also known as superfunctionalization [37] –, thereby adding robustness to mammalian regulatory networks.

On average, a shadow enhancer pair had common activity in 1.2 facets and 1,328 pairs (63%) had 100% identical activity profiles (see Figure 1E). Furthermore, consistent with previous studies [18,32] and supporting the idea that regulatory redundancy is scattered across more than two enhancers, we found that 672 shadow enhancers (52%) were partners of multiple shadow enhancer pairs. To gain insight into such complex relationships, we arranged shadow enhancers into groups, such that all enhancers in a group have the same target genes and common activity with at least one other enhancer. On average, each group consisted of 2.7 enhancers (see Figure 1F). Two shadow enhancer groups consisting of 34 and 21 enhancers were exceptionally large. An explanation for such a high number of redundant enhancers could be a very complex spatio-temporal expression profile of the targets in these groups. The group of 35 enhancers was associated with eight transcripts of the *thyroglobulin* (ENSG00000042832) gene, which is mainly expressed in the thyroid gland (see Methods). Interestingly, all enhancers in this group have highly redundant regulatory activities, with all of them being active in the “thyroid” facet as well. This suggests that *thyroglobulin* is transcribed in a complex condition-specific manner, rather than having a complex spatial expression pattern. Similarly, the group of 21 enhancers apparently regulates the expression of three transcripts of the *ADAM metallopeptidase domain 12* (ENSG00000148848) gene in the placenta. In addition to demonstrating differences in sizes, shadow enhancer groups differed in the number of associated target transcripts: while 47% of the groups had only one target transcript, 53% had multiple ones; the average number of associated target transcripts over all groups was 2.3. This result is supported by other genetic and genome-wide studies [38,39] and indicates a relatively high level of regulatory complexity, with multiple enhancers being associated with multiple promoters. Lastly, although not every pair of enhancers in a group was required to have common activity, almost all of them did (2,072 out of a total of 2,092 pairs). Thus, the majority of shadow enhancer pairs in our dataset shared activity in one facet and were not active in any other facet, showing a very high level of redundancy.

### A large fraction of human shadow enhancer pairs have independent origins

Shadow enhancers have been proposed to arise by duplication [5]. Nevertheless, this hypothesis has not been systematically tested and remains speculative. Indeed, it has been clearly established that a large number of regulatory elements derived from transposons [40], and examples of independent exaptation of transposons into shadow enhancers are known [27]. In order to quantify the contribution of transposons to the genesis and evolution of shadow enhancers, we annotated each enhancer based on its overlap with RepeatMasker elements (see Methods). Out of the 3,523 enhancers that were correlated with at least one promoter, 1,686 (48%) were annotated as transposons and, hence, may derive from a transposon. We refer to these enhancers as transposon enhancers. In agreement with previous observations [41], this number is lower than expected from their prevalence in the genome (log2-fold difference in observed transposon overlap compared with the random expectation = −0.5, see Figure S2). Furthermore, also in accordance with other studies [42,43], the fraction of enhancers annotated as transposons depends on the facet, and varies from 24% (adipose) to 66% (fingernail, see Figure S3). While most of the transposon enhancers (1,132 out of 1,686, 67%) were annotated as transposons from only one transposon species (see Methods), 554 (32.9%) were annotated with multiple (two to four) transposon species, and may include cases of coordinate co-option of transposons [23]. The five most represented transposon families among the enhancers were L2s (20.7%), Alus (20.1%), MIRs (17.6%), L1s (12.6%) and ERVL-MaLRs (5.4%, see Figure 2A). These are also the most prominent transposon families in the human genome. However, whereas L2s and MIRs were over-represented among the enhancers as compared to their frequencies in the genome, L1s, Alus and ERVL-MALRs were depleted (empirical P-values < 0.05, see Methods). The fraction of enhancers annotated as transposons did not differ significantly between shadow and non-shadow enhancers. Moreover, we found no differences between shadow and non-shadow enhancers when comparing the fraction of enhancers overlapping with a particular transposon family. This suggests that transposons have been similarly co-opted as shadow and non-shadow enhancers.

**Figure 2.**
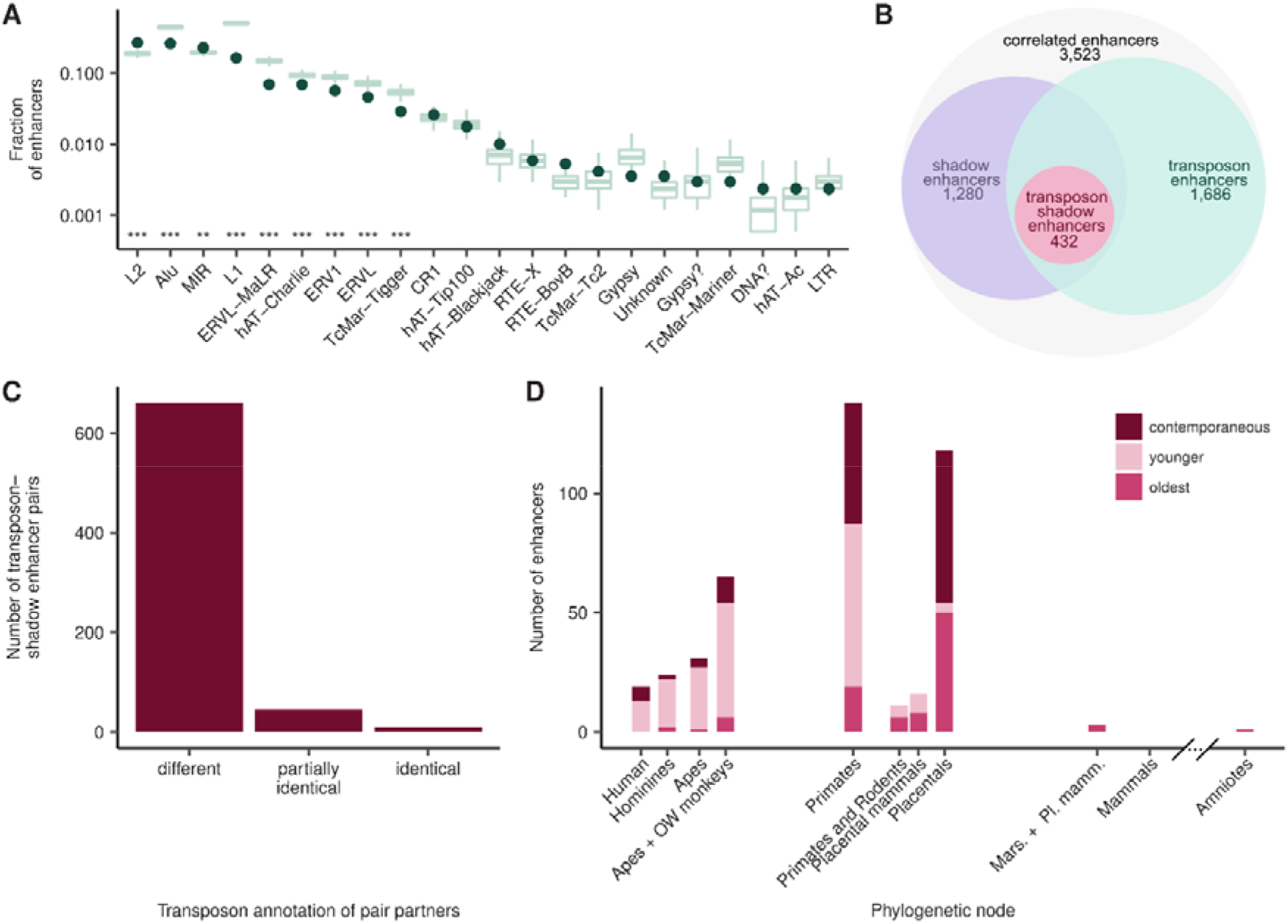
Most partners of transposon-shadow enhancer pairs originated independently. **A)** Transposon families contribute differently to the evolution of shadow enhancers. The green dots indicate the fraction of transposon enhancers annotated with each transposon family. The green boxplots depict the distribution of the fraction of random genomic sequences annotated with the same transposon family, estimated based on 1000 random sets. Asterisks indicate adjusted empirical P-values. ’*’ P < 0.05, ’**’ P < 0.01, ’***’ P < 0.001. **B)** The scheme depicts the connection of the shadow enhancer, transposon enhancer and transposon-shadow enhancer sets. The 3,523 enhancers that are significantly correlated to a FANTOM promoter are referred to as “correlated enhancers”. 1,686 correlated enhancers annotated as transposons and are referred to as “transposon enhancers”. 1,280 correlated enhancers form one or more shadow enhancer pairs and are referred to as “shadow enhancers”. Finally, if both partners of a shadow enhancer pair are annotated as transposons, they are referred to as “transposon-shadow enhancers”; these pairs comprise a total of 432 enhancers. **C)** In the majority of transposon-shadow enhancer pairs the partners have different transposon annotation. Depicted are the number of pairs where partners have different, partially identical (i.e., some transposons are identical) or identical transposon species annotation. **D)** Redundancy in transposon-shadow enhancers seems to have originated mostly during the evolution of placentals. Shown is the age of transposon-shadow enhancers, estimated based on the insertion times of the transposons with which they are annotated. Redundancy can arise by i) the evolution of a new (“younger”) enhancer with regulatory activities that are (possibly partially) redundant to those of an already existing (“oldest”) enhancer; or ii) by the simultaneous (“contemporaneous”) evolution of two or more enhancers with (possibly partially) regulatory activities.

Out of the 2,072 shadow enhancer pairs, 716 (35%) were transposon-shadow enhancer pairs, in the sense both enhancers of the pair overlap with transposons. These transposon-shadow enhancer pairs involved a total of 432 enhancers, to which we further refer as transposon-shadow enhancers (see Table 1 and Figure 2B). The number of transposon-shadow enhancer pairs is consistent with the expectation based on the tissue-specific activities of our shadow enhancers and the rates of transposon exaptations into enhancers that have been reported in the past [41]. Remarkably, the partners of the vast majority (661, 92%) of transposon-shadow enhancer pairs had different transposon species annotation (see Figure 2C), indicating that the partners mostly derive from independent transposon insertions. In order to account for possible inaccuracies in the transposon annotation, we also compared the annotation of the enhancers on the family level of the transposon taxonomy (see Methods). Although the number of transposon-shadow enhancer pairs with different transposon annotation was smaller (466, 65%), it was still the majority. Moreover, increasing the stringency to annotate enhancers as transposon enhancers (see Methods) resulted in at most a moderate decrease in the fraction of transposon-shadow enhancer pairs with different transposon annotation, both on the species and family levels of the transposon annotation (see Figure S4). These results suggest that in the majority of transposon-shadow enhancer pairs the two enhancers have independent origins.

**Table 1.** Number of enhancers, enhancer pairs and enhancer groups in the human genome, for multiple sets. The 3,523 enhancers that are significantly correlated to a FANTOM promoter are referred to as “correlated enhancers”. 1,686 correlated enhancers overlap with transposons and are referred to as “transposon enhancers”. 1,280 correlated enhancers form one or more shadow enhancer pairs and are referred to as “shadow enhancers”. Finally, if both partners of a shadow enhancer pair are annotated as transposons, they are referred to as “transposon- shadow enhancers”; these pairs comprise a total of 432 enhancers. Analogously to shadow enhancer groups, transposon- shadow enhancer pairs can also be arranged into transposon-shadow enhancer groups.

| Set | Nr of enhancers | Nr of pairs | Nr of groups | Coloring in main figures |
| --- | --- | --- | --- | --- |
| Correlated enhancers | 3,523 | - | - | Gray |
| Shadow enhancers | 1,280 | 2,117 | 465 | Purple |
| Transposon enhancers | 1,686 | - | - | Green |
| Transposon-shadow enhancers | 432 | 716 | 157 | Red/raspberry |

To obtain a more comprehensive picture of the origin of our transposon-shadow enhancers, we inferred the evolutionary age of the corresponding transposons using a maximum parsimony approach (see Methods). Specifically, we first searched for candidate orthologous sequences for the transposon-shadow enhancers in the genomes of 18 mammalian and vertebrate species using BLAST (see Figure S5). Next, we required the candidate orthologues to be located in the neighborhood of a target gene orthologue and to overlap with the same transposon types as the human enhancer. Thus, from our transposon-shadow enhancers 96% had an orthologue among other primate species, 22% an orthologue in rodents, 47% in the Laurasiatheria and 1% in the opossum branch. In general, as anticipated, the less a clade was related to human, the fewer enhancers had an orthologue in the clade. The only exception was the rodent clade, which had relatively few orthologues despite the close relationship. This can be explained by the fact that rodents have a faster molecular clock than primates [44]. Subsequent phylogenetic reconstruction (see Methods) showed that most of the transposon insertions in our transposon-shadow enhancer data set date back to the common ancestor of placentals (25%, 105.5 MYA) or primates (32%, 73.8 MYA), and this is roughly in accordance with the estimated activity periods of the transposons. However, compared to the overall age distribution of transposons in the human genome, the transposons in the transposon-shadow enhancers were older (mean ages: 69.16 MYA vs. 76.0 MYA; P-value = 6.26 × 10^−4^, two-sided Wilcoxon rank-sum test, see Figure S6 and Methods). This is in agreement with previous reports that enhancers are enriched in ancient transposons [41]. In particular, we observed that 3% (4) of the transposon-shadow enhancers derived from a transposon acquired in the common ancestor of placentals were redundant to others originating earlier, while 42% (50) possibly represent cases in which redundancy evolved in a contemporaneous manner (see Figure 2D). Hence, the transposon-derived regulatory network in the common ancestor of placentals appears to have had a level of redundancy of at least 45%. This is even more prominent in the primate ancestor. Indeed, 49% (68 enhancers) of enhancers originating in the common primate ancestor were redundant to enhancers with older transposons and 37% (51 enhancers) evolved contemporaneously. Interestingly, we found that among transposon-shadow enhancer pairs in which at least one of the partners derived from a transposon insertion dating to the common primate ancestor or it successors – but only among them –, the younger partners had a significantly higher SNP density than their older counterparts (medians 116 vs 106 SNPs/kb, adjusted P-value = 3.6×10^−5^). This suggests that the younger partners of those pairs are less constrained to evolve than the older partners. Overall, the median absolute age difference between the partners of transposon-shadow enhancer pairs was 31.7 MY, suggesting that redundancy is often introduced into regulatory networks after long periods of time. Together, these findings point to the introduction of redundancy via independent transposon insertions as a common event in mammalian evolution.

To investigate to which degree the introduction of regulatory redundancy is driven by the acquisition of new genes, we estimated the age of the transposon-shadow enhancer target genes and compared it to that of the corresponding transposons (see Methods). The majority of the genes (2,473 out of 2,976 target genes, see Figure S7) date back to the “bony vertebrates” or older clades and are, thus, presumably to a great extent older than the enhancers in our dataset that are associated with them. This already indicates that newly originating genes are not a main driver for the introduction of regulatory redundancy in our data. Only seven transposon-shadow enhancer pairs had gene targets that were younger than one of the partners in the pair, and among those, there were five – all belonging to the same shadow enhancer group – for which one of the target genes originated after the older enhancer, but before the younger one: DRICH1, with expression in many tissues and a high expression in testis [45]. Hence, with a few exceptions, the emergence and fixation of regulatory redundancy is not associated with the acquisition of new genes. Instead, our evidence points towards gene superfunctionalization [37].

### Regulatory redundancy is mostly lineage-specific

In order to establish to what extent our finding of a widespread independent origin of shadow enhancers holds true for the genomes of other species, we conducted the same analyses for mouse (see Methods). From originally 44,459 mouse FANTOM enhancers and 158,965 promoters we identified 4,074 enhancers with a significant correlation coefficient to 8,082 promoters (see Tables S1 and S2). Among those, we distinguished 1,939 shadow enhancers, forming 2,787 shadow enhancer pairs (see Figures 3A and B, Table S3). In contrast to their human counterparts, mouse shadow enhancers did not differ in their SNP-density from non-shadow enhancers, but they had a slightly lower GC-content (with medians of 0.50 and 0.49, respectively, P-value = 3.4 × 10^−6^, two-sided Wilcoxon rank-sum test). Analogously to what we observed in the human genome, the target genes of shadow enhancers in the mouse genome were more tissue-specific (median Shannon entropy of 2.5 for non-shadow enhancer target genes and 2.3 for shadow enhancer target genes, P-value = 1.1×10^−3^, two-sided Wilcoxon rank-sum test, see Methods and Figure S8) and tended to be more strongly expressed (in 53% of the evaluated facets, with no difference in the remaining 47%, adjusted P-values ≤ 0.05, see Methods) than those of non-shadow enhancers. We further observed that mouse shadow enhancers were, like human shadow enhancers, over-represented in thymus (adjusted P-value = 1.2×10^−9^, see Methods) and depleted in brain-related facets such as spinal cord (P-value = 1.1×10^−21^), brain (P-value = 7.8×10^−15^) and pituitary gland (P-value = 2.8×10^−10^). Mouse shadow enhancer pairs also showed a very high level of redundancy, sharing common activity in an average of 1.1 facets. Moreover, 1,961 pairs (70%) had 100% identical activity profiles. Taken together, these results indicate that mouse and human shadow enhancers fulfill similar functions, despite differences in their evolutionary divergence and genomic location.

**Figure 3.**
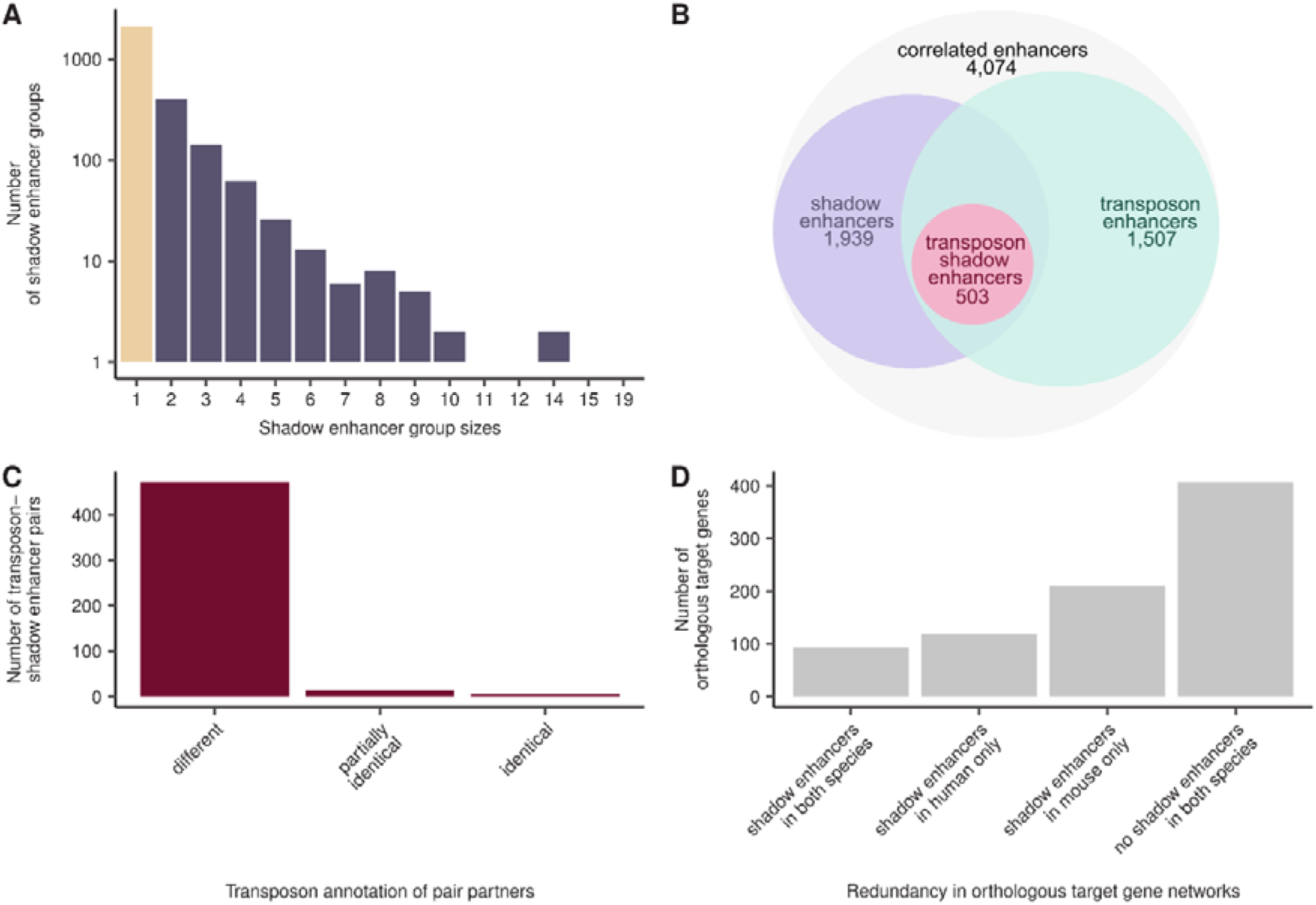
Shadow enhancers in the mouse genome show similar properties to those in the human genome, but redundancy in orthologous regulatory networks is mainly lineage-specific. **A)** Shadow enhancers in the mouse genome form groups of different sizes, similar as in human. Depicted are the number of non-shadow enhancers (beige) and shadow-enhancer groups (purple). **B)** The scheme depicts the connection of the shadow enhancer, transposon enhancer and transposon-shadow enhancer sets. See Figure 2 B) for details. **C)** The enhancers of shadow enhancer pairs show the same transposon annotation pattern as in human. Depicted are the number of transposon-shadow enhancer pairs where partners have different, partially identical (some transposons are identical) or identical transposon species annotation. **D)** A large fraction of target genes that are orthologous between human and mouse are regulated by shadow enhancers in just one of the two species. Shown 25 are the common orthologous enhancer target genes between the human and mouse correlated enhancer datasets and their regulation by shadow enhancers in human and mouse.

Compared to human, a substantially smaller fraction of mouse enhancers were transposon enhancers (37%, log2-fold difference in observed transposon overlap compared with the random expectation = −0.77, see Figure S9). This may be partially explained by the faster molecular clock of mouse [44], and the ensuing difficulty to recognize transposon-derived sequences as such. The five most prominent transposon families among mouse enhancers were the Alu (17% of transposons), B4 (16%), ERVL-MaLR (12%), L1 (11%) and ERVK (8%) families (see Figure S10). Thus, while Alus were the most prominent family in both human and mouse, the ERVK family was much more frequent in mouse and MIRs and L2s were less frequent (7% and 4%, ranking in 7^th^ and 8^th^ position, respectively, after B2 with 7%) in mouse than they were in human. As observed for human, L2s and MIRs were over-represented among the enhancers as compared to their frequencies in the genome, while Alus, L1s, ERVKs and ERVL-MaLRs were depleted (empirical P-values ≤ 0.05, see Methods). Finally, although the rodent specific family B4 made up a large fraction of the mouse enhancers, its frequency matched the genomic expectation. Hence, many transposon families display the same trend towards exaptation into enhancers in both human and mouse. Further, we did not find any difference between the transposon families represented among shadow and non-shadow enhancers. Reflecting the lower number of transposon-derived enhancers in mouse, only 18% (493) of shadow enhancer pairs were transposon-shadow enhancer pairs, involving 503 transposon-shadow enhancers. Nevertheless, similarly to what we observed in the human genome, the majority of them had different transposon species (96%, see Figure 3C) and family (76%) annotations, providing further support for the hypothesis that transposon-shadow enhancers in mammalian genomes mainly originate by independent exaptation events. Interestingly, only few (17%) transposon-shadow enhancers in mouse had an orthologue in other mammalian species, mainly among primates and Laurasiatheria (15% and 13%, respectively); only 6% had an orthologue in rabbit, the only other member of the glires considered in the analysis. Consequently, 93% (457) of transposon-shadow enhancers – and, in general, a vast fraction of enhancers – appeared to be mouse specific. Although this high fraction could be partly explained by the fact that the species included in the phylogenetic analysis were only distantly related to mouse (see Methods, Figure S5), the fraction of transposon-shadow enhancers derived from transposons dating back to the common ancestor of placentals and rodents or before was smaller in mouse (13%) as compared to human (35%), indicating that currently active transposon-shadow enhancers are younger in mouse than in human. In summary, the transposon-shadow enhancers in mouse are composed of other transposon families than in human, but still the vast majority (96%) of transposon-shadow enhancer pairs consists of partners that are apparently derived from different transposon species (76% on transposon family level) – and thus 17% (13% on transposon family level) of all shadow enhancer pairs in the mouse genome appear to have evolved independently and in parallel.

Out of 828 enhancer target genes that were common between the human and mouse enhancer datasets (see Methods), only 93 (11%) were regulated by shadow enhancers in both species, whereas 328 (39%) were regulated by shadow enhancers in one species, but not in the other (see Figure 3D). The number of genes regulated by shadow enhancers in both species is small, but greater than expected by chance (odds-ratio = 1.5, P-value = 6.0 × 10^−3^, one-sided Fisher’s exact test), which hints at a requirement of redundancy in the regulatory networks of both species. However, shadow enhancers appear to be mainly species specific. Indeed, only 36 (1%) human enhancers had an orthologue in mouse (see Methods), and only nine (2%) out the 556 human shadow enhancers with orthologous target genes in mouse were orthologous to the corresponding mouse shadow enhancers. Conversely, only eleven (0.3%) mouse enhancers had an orthologue in human, and only four out of the 880 mouse shadow enhancers with orthologous target genes had orthologues in human. This indicates that the enhancers driving the expression of the genes that are regulated in a redundant manner in both human and mouse have evolved independently. Together, our results suggest that, with some exceptions, evolution of regulatory redundancy is a largely species-specific process among mammals, both concerning the target genes and the enhancers, and that orthologous regulatory networks with similar levels of redundancy are likely to result from convergent evolutionary processes.

## Discussion

It has long been hypothesized that shadow enhancers originate by duplication [5]. However, exaptation or co-option of transposons has been often reported as source of enhancers in the mammalian genome [26,46,47], also of shadow enhancers [27]. Our study shows that a large number of shadow enhancers indeed appear to have arisen from independent transposon exaptations.

We chose to base our analysis on an enhancer dataset generated by the FANTOM5 consortium using CAGE. Thus, our results are conceivably limited to enhancers that produce bi-directional eRNA. Whether this is a property that every active enhancer shows, is still a matter of debate. However, it is likely that not all active enhancers have this property [48]. In any case, the features distinguishing these alleged two classes of enhancers remain unclear, and therefore, it is difficult to assess to what degree our findings could be extended to enhancers that are not transcribed. Importantly, the dataset has a high resolution (enhancer activity is quantified at a single base pair resolution) and contains a large number of tissue samples, which allowed for a target gene assignment via correlation. Correlation is a frequently used approach and we used a correlation-based target gene assignment strategy similar to the one in the FANTOM5 enhancer study [31,49–51]. Enhancer-promoter interactions are known to mainly occur within TADs [52]. Thus, we only tested enhancer-promoter pairs where both partners are located within the same TAD for correlation, as opposed to testing within windows of a fixed size. Moreover, we required shadow enhancers to be associated with the exact same set of target transcripts as opposed to only at least one of them. This leads to a slightly smaller number of shadow enhancer pairs than when pairing all enhancers with at least one common target transcript, but makes the pairing more conservative. The number of shadow enhancer pairs active in a given facet varied greatly, as did the total number of enhancers active in that facet. Accordingly, many of our findings are expected to reflect the tissue-specific activities of the shadow enhancers in our dataset. Finally, although our shadow enhancer pairs are of a putative nature, they rely on data that have been extensively validated over the past five years. Indeed, the enhancer predictions in the FANTOM5 dataset have been: i) shown to overlap by at least 71% with predictions based on DNase-I-Hypersensitivity, H3K27ac andH3K4me1 histone modifications [31]; ii) successfully tested in zebrafish reporter gene assays [31]; scrutinized to demonstrate other enhancer attributes (e.g., [36,53]).

Our results indicate that shadow enhancers are common features of regulatory networks not only in *Drosophila* [18], but also in mammals. In contrast to some of previous studies [18,27,54], we found that the majority of our shadow enhancers were completely redundant in terms of their expression pattern. However, this result depends on the number of tissue samples available. While the number of tissues that was involved in the current analysis is relatively large, our shadow enhancers may still diverge in their activity in other tissues or under different conditions. In addition, some of our facets comprise several distinct cell types, which could lead to an overestimation of the number of (completely) redundant enhancers. It was further hypothesized before that the partners of highly redundant enhancer pairs might be subjected to different selective pressures, with one enhancer being less conserved than the other and free to evolve new – ultimately non-redundant – functions. Although this is supported by the slightly higher SNP densities that we noted for relatively young enhancer partners, it is not consistent with the fact that shadow enhancers were indistinguishable from non-shadow enhancers – their non-redundant counterparts – in terms of sequence conservation, which has also been observed in *Drosophila* [18]. On the contrary, our data suggest that regulatory redundancy provides a selective advantage that would contribute to the fixation of shadow enhancers. Indeed, in agreement with previous findings [14,55,56], we showed that shadow enhancers are associated with more precise and higher target gene expression.

Approximately 50% of the human (and 40% of the mouse) shadow enhancers were annotated as transposons and, as such, represent possible cases of transposon exaptation. In agreement with observations for non-shadow enhancers [41,42], our shadow enhancers were generally depleted of transposons compared to the entire genome. This does not imply that the number of enhancers deriving from transposons is small; it is just smaller than expected from the fraction of the total genome that is derived from transposons. Several factors could account for this. Specifically, given the low sequence conservation of many enhancers, a fair amount of enhancers might be derived from ancient transposons for which the transposon signature is not visible anymore. Furthermore, it appears plausible that only some transposon families are well suited to be co-opted into enhancers. Certainly, our enhancers were only enriched among particular transposon families. In addition, transposons carry TFBS that lead to activity in certain tissues and, therefore, the different enrichment levels could be partially explained by the tissue-specificities of the enhancers represented in the dataset. Interestingly, the vast majority of our transposon-shadow enhancer pairs in both human and mouse comprised enhancers that were most presumably derived from different transposon species, and are, thus, likely to have evolved from independent transposon exaptation events. Increasing the stringency in the identification of enhancers annotated as transposons led to only a mild decrease in the number of transposon-shadow enhancer pairs involving different transposon species, if any.

Our phylogenetic analysis suggests that most transposon insertions associated with the origination of human shadow enhancers took place in the primate or placental ancestors, and that in at least 77% of transposon-shadow enhancer pairs the partners originated several tens of millions of years after one another. In the remaining 23% of the cases the partners in the pairs might have evolved in a contemporaneous manner. However, it must be noted that the primate and placental common ancestor nodes in our phylogenetic tree cover relatively large periods. Consequently, what we consider contemporaneous, could actually be several tens of MY apart. Unfortunately, this cannot be completely circumvented since there are no recent species that would allow to estimate the ancestral state for certain periods. Furthermore, this analysis disregards redundancy introduced from non-transposon shadow enhancers. As some non-transposon enhancers likely represent ancient transposon-derived enhancers, redundancy is likely to have been first introduced earlier than the estimated date. The mouse transposon-shadow enhancers were mostly mouse-specific and, thus, substantially younger than those in human. This is expected due to the faster evolution of the mouse genome, but is also impacted by the degree of relatedness of the species included in the phylogenetic analysis. In addition, the quality of the genome assembly of the most closely related species (rabbit) is not as high as the assemblies of the primate species, and we may have failed to detect some orthologues in rabbit due to gaps in its assembly. In any case, even if the transposon insertions had been contemporaneous, there could still be millions of years between the transposon insertions and their exaptation into enhancers. Finally, many of the shadow enhancers that were not annotated as transposons and, hence, disregarded in these calculations, could actually be ancient transposon-derived sequences for which the transposon signature is not traceable anymore. Therefore, our results presumably underestimate the extent to which independent transposon exaptation has contributed to shadow enhancer origination.

We found hardly any orthologues among human and mouse shadow enhancers, which is consistent with rapid enhancer turnover [10,43,57–59]. Nevertheless, we observed that among orthologous target genes the number of genes with shadow enhancers in both species was larger than expected by chance. Since the corresponding shadow enhancer pairs were not orthologous, the redundancy must have developed independently. Hence, certain regulatory networks apparently have a higher propensity – or at least a higher tolerance – towards regulatory redundancy. Given the properties that are associated with regulatory redundancy – like a higher target gene expression – we hypothesize that these networks benefit from redundancy. Nevertheless, most orthologous regulatory networks showed differential redundancy, in the sense not all the genes that were redundantly regulated in one species were redundantly regulated in the other and, in the cases in which they were, the regulation was actualized by non-orthologous *cis*-regulatory elements. In the light of the enhancer turnover model and the low sequence conservation of many known enhancers [58], regulatory redundancy in some of these networks may be merely temporal and not exerting an essential function. Yet, the clear association of shadow enhancers with gene expression strength and tissue specificity suggests that regulatory redundancy *per se* is an important property of regulatory networks.

In summary, the redundant partners in 92% of all transposon-shadow enhancer pairs – 31% of all shadow enhancer pairs – in the human genome have been acquired independently, in successive waves of transposon expansions. The mouse genome displays comparable numbers (95% of mouse transposon-shadow enhancer pairs – 17% of all shadow enhancer pairs – have independent origins). Shadow enhancers were previously hypothesized to derive from sequence duplications. Here, we show that independent transposon exaptation constitutes an alternative mechanism. Considering that many enhancers without a transposon signature are likely to derive from ancient transposons, this may even be the predominant mechanism of shadow enhancer origination. Moreover, despite shadow enhancers being a common feature of many mammalian regulatory networks, shadow enhancers are poorly conserved and mostly lineage-specific, with hardly any orthologous shadow-enhancers between different lineages. Hence, most regulatory redundancy in mammalian networks appears to have developed independently by convergent evolution. In any event, given its widespread distribution, understanding the evolutionary processes by which regulatory redundancy arises is key to understanding how networks respond to perturbations, in particular, those associated with disease.

## Materials And Methods

### Multiple hypothesis testing adjustments

P-values were adjusted for multiple testing by controlling the false discovery rate [61].

### Genome annotation

Annotation was based on GENCODE (human: v19, mouse: vM1, [62]).

### FANTOM data and processing of hCAGE read counts

We used cell type specific enhancer and promoter activity data from the FANTOM5 Project for human (hg19) and mouse (mm9), phase 2.5 [30,31]. Specifically, we used the hCAGE data for the samples that were labelled organ derived for human and mouse (http://fantom.gsc.riken.jp/5/datafiles/phase2.5/basic/human.tissue.hCAGE/ and http://fantom.gsc.riken.jp/5/datafiles/phase2.5/basic/mouse.tissue.hCAGE/). We further used the sets of promoter and enhancer coordinates from FANTOM (http://fantom.gsc.riken.jp/5/datafiles/phase2.5/extra/Enhancers/ and http://fantom.gsc.riken.jp/5/datafiles/phase2.5/extra/CAGE_peaks/).

### Processing of hCAGE read counts

Reads with MAPQ < 20 were filtered out.

### Grouping and merging of tissue samples into facets

We grouped samples from similar tissues into “facets” (we used the same assignment of samples to facets as Andersson et al., [31]). We then merged the hCAGE bam files of the sample groups into “facets”. This resulted in 44 tissue facets for human and 36 tissue facets for mouse.

### Computation of raw counts for enhancers and promoters

We computed the read counts for enhancers according to [31] and for promoters according to [30]. We took a 400 bp window around the midpoint of the enhancers, removing all enhancers that overlapped with any FANTOM promoter or with ENSEMBL gene exons. For both enhancers and promoters we counted all hCAGE reads where the 3’ end overlapped the enhancer (resp. promoter), that had an edit distance ≤ 6 (NM flag) and a MAPQ ≥ 20.

### Computation of tags per million reads (TPM) for enhancers and promoters

We computed CAGE tags per million mapped reads according to FANTOM [31]: We defined the raw library size as the total number of mapped reads after MAPQ filtering, without reads on chromosome M, and computed library size normalization factors with the edgeR package function calcNormFactors() and its “RLE” method.

### Facet specific binary enhancer and promoter activity (“enhancer/promoter usage matrix”)

To compute binary enhancer activity (active/inactive) we followed the rationale and approach of Andersson et al. [31]. We sampled 100,000 401 bp-long regions from the human genome (resp. mouse). The regions were ensured to not overlap with ENSEMBL exons, FANTOM promoters or FANTOM enhancers. We counted reads in these random regions in the same manner as for the FANTOM enhancers/promoters and computed an empirical P-value as the fraction of random sequences with equal or greater raw read counts in comparison to the enhancers and corrected for multiple testing. We considered an enhancer active in a specific facet if it had at least 2 reads, a P-value ≤ 0.0025 and an FDR-corrected P-value ≤ 0.1.

For the promoters, we computed the facet specific binary promoter activity (“promoter usage matrix”) according to FANTOM. Andersson et al. used a cutoff of 0.2 on the normalized activity values [31]. We computed the corresponding percentile and used the value of the corresponding percentile as the threshold for our data (human: 0.297, mouse: 0.319).

### Correlation of facet-specific promoter and enhancer activities and formation of shadow enhancer groups

We only considered enhancers and promoters active in at least one of the facets used in this study (see “Facet specific binary enhancer and promoter activity (“enhancer/promoter usage matrix”)”). We assigned all ENSEMBL TSS of protein coding genes with a distance of ≤ 500 bp to the corresponding promoters and discarded promoters without a TSS assignment. We further assigned all enhancers and promoters to a TAD if their sequence overlapped with it at least 50% and discarded enhancers and promoters not assigned to any TAD. We then mapped the enhancer and promoter TPM values to a simplified activity scale: Enhancers (promoters) that were inactive in a tissue according to the usage matrix were assigned a value of zero, the rest was split into tertiles across all facets and their activity values replaced by 1, 2 or 3. Finally, we computed the Pearson correlation coefficient for each pair of enhancers and promoters assigned to the same TAD based on their simplified activities, and tested for association between the enhancer and promoter activities using the Pearson product-moment correlation test. P-values were FDR-corrected across all tested associations. We considered all enhancer-promoter associations with r ≥ 0.5 a P-value ≤ 0.0035 and an FDR-corrected P-value of ≤ 0.05 as significant.

### Tissue specificity of target genes

We computed the Shannon entropy [63] for each target gene. The computation was based on the TPM values across all facets of all its correlated (FANTOM) promoters. If a target gene was associated with multiple FANTOM promoters, we averaged the entropy across all promoters. We compared the entropy of non-shadow enhancer target genes to shadow enhancer target genes using a two-sided Wilcoxon rank-sum test. Each target gene was considered only once and those in both groups were excluded.

### Target gene expression strength

For each facet, we measured the expression strength of a gene as the average of the TPM values of all its promoters. Genes with multiple promoters active in that facet such that at least one of the promoters was associated with a shadow enhancer and at least one of the promoters was associated with a non-shadow enhancer were excluded from the analysis. Then, we compared the expression strength between shadow enhancer target genes (i.e., genes with two or more active enhancers in that facet) and non-shadow enhancer target genes (i.e., genes with only one active enhancer in that facet) with a one-sided Wilcoxon rank sum test. P-values were corrected for multiple testing. Only facets with more than ten shadow and non-shadow enhancer target genes were considered.

### Shared activity of a shadow enhancer pair

To measure the degree of shared activity of a shadow enhancer pair, we computed the fraction of facets with activity of both enhancers (intersection of active facets) and tissues with activity in either enhancer (union of active facets). The enhancers were regarded as active or inactive in a given facet according to the usage matrix.

### Sequence conservation of enhancers

Per enhancer we computed a total sequence conservation score as the average of the base-wise PhastCons scores [64]. For human the computation was based on the hg19 100way phastCons scores from UCSC (http://hgdownload.cse.ucsc.edu/goldenpath/hg19/phastCons100way/hg19.100way.phastCons.bw) and for mouse on the mm9 30way placental phastCons scores (http://hgdownload.cse.ucsc.edu/goldenpath/mm9/phastCons30way/placental/).

### SNP density of enhancers

We computed the SNP density as SNPs per 1000 bp with data from ENSEMBL variation [65] (ftp://ftp.ensembl.org/pub/release-92/variation/vcf/homo_sapiens/homo_sapiens.vcf.gz, ftp://ftp.ensembl.org/pub/release-92/variation/vcf/mus_musculus/mus_musculus.vcf.gz). The enhancer coordinates were lifted over to hg38 and mm10. Enhancers that could not be lifted over were discarded from this analysis.

### Facet enhancer enrichment

For each facet we compared the number of active and inactive enhancers between shadow and non-shadow enhancers. We computed a P-value with a two sided Fisher’s exact test and corrected for multiple testing.

### Target gene expression

Target gene expression patterns were compared to the ENSEMBL expression atlas [45].

### Transposon annotation of enhancers

Enhancer sequences overlapping by at least 50 bp with one or more transposons of the same species (possibly interrupted by an arbitrary sequence) in the RepeatMasker database (http://www.repeatmasker.org, version 4.0.5) were annotated as transposons. For enhancers shorter than 500 bp, we used a 500 bp window around the center point of the original coordinates. The RepeatMasker taxonomy classifies repeats (“species”) into “families” which are, in turn, classified into “classes” (“LINE”, “SINE”, “LTR”, “DNA”, etc). Only the repeat classes “LINE”, “SINE”, “LTR” and “DNA” were used in the analysis. Enhancers satisfying the overlap requirement for multiple transposon species were annotated with all of them.

### Genomic distribution of transposons

We modelled the genomic transposon overlap by randomly repositioning our 3,523 correlated enhancers (4,074 in mouse) in the genome and computing the transposon overlap in the same was as for the enhancers. This was repeated 1000 times and the derived background distributions were used to test every transposon family for significant enrichment among our shadow enhancers and to compute an empirical P-value.

### Enrichment of transposon families among shadow and non-shadow enhancers

For every transposon family, we counted the number of shadow and non-shadow enhancers annotated as transposons of that family. Then, we tested the null hypothesis of equal proportions (two-sided test) among shadow and non-shadow enhancers in R with the prop.test() function.

### Pairwise comparisons of shadow enhancer transposon annotation

We counted the number of transposon-shadow enhancer pairs in which the partners were annotated with different, identical or partially identical (in case of multi-transposon enhancers) transposon species (or families). We repeated this with increased stringency in the transposon annotation, considering only those pairs where both partners showed a minimum total overlap with one or more transposons of 10% to 100% of their sequence, in 10% steps.

### Identification of transposon-derived enhancer orthologues

We searched for possible orthologous sequences of the human FANTOM enhancer sequences in a number of mammalian and vertebrate species. The assemblies we used were panTro4, gorGor3, ponAbe2, rheMac8, calJac3, mm9, oryCun2, bosTau8, canFam3, myoLuc2, loxAfr3, dasNov3, monDom5, ornAna2, galGal4, xenTro7, fr3 and danRer10 as provided by the UCSC genome browser website. The species and assembly versions were selected based on the quality of available assemblies, availability of ENSEMBL Compara data for the assemblies and in such a way as to sample all big branches of the vertebrate phylogenetic tree with focus on mammals and primates [66]. We used the same windows around enhancers shorter than 500 bp as for the transposon classification. We employed blastn from the NCBI BLAST+ suite (version 2.2.31, [67]) selecting scoring parameters that promote the score of sequences with moderate similarity (target sequence similarity ~ 70%) [68]: word size 7, reward 5, penalty −4, gap opening cost 8, gap extension cost 6 and an E-value cutoff of 1×10^−3^. In addition to being one of the ten highest scoring BLAST matches the enhancers had to fulfill the following requirements in order to be labelled the orthologous sequence. For every enhancer we identified the orthologues of all its human target genes via ENSEMBL Compara (release 87, hg38). All gene orthology relationships (one2one, one2many, many2many) were included. The ENSEMBL hg38 gene IDs were mapped to hg19 gene IDs (as some IDs are not stable) and the gene orthologue coordinates of some species had to be lifted over to the used assembly. We required the BLAST matches to be located within a 2 Mb window around any target gene orthologue. Enhancers without any target gene orthologues were excluded from the analysis. In addition, we required the BLAST matches of human transposon enhancers to overlap with the same RepeatMasker transposon types and number of transposons as the human enhancer (RepeatMasker version 4.0.5; mm9: lift over from mm10, bosTau8: 4.0.5, rheMac8: 4.0.5, ornAna2: 4.0.5). In order to reduce the effect of slight annotation differences across the vertebrate genomes we considered the transposon types identical to human if they belonged to the same transposon family. For enhancers overlapping with multiple transposons this meant all transposon families had to be there in order for the orthologue to be called in the assembly. In case of multiple BLAST matches fulfilling these criteria, we declared the match with the highest E-value the enhancer orthologue.

### Phylogenetic reconstruction

The tree topology for the selected vertebrate species (see “Identification of transposon-derived enhancer orthologues”) was generated with phyloT (http://phylot.biobyte.de/) with a random breakdown of polytomies. Using binary states that represented the presence or absence of the enhancer orthologue, we then reconstructed the ancestral states in the tree for every enhancer with the phangorn R package. After importing the tree topology with the read.newick() function (phytools package), deleting single nodes and setting all branch lengths to one with the collapse.singles() and compute.brlen() functions (ape package), respectively, we used the ancestral.pars() function (phangorn package) with type=MPR for a maximum parsimony reconstruction (Fitch-Hartigan algorithm). We defined the insertion node as the oldest node in the human (or mouse) lineage with an uninterrupted sequence of nodes (starting at the human/mouse node), where the orthologue was inferred to be present. For a few enhancers without a target gene orthologue in any of the used vertebrate species, the dating was not possible. For the enhancers overlapping with multiple transposons, the age of the youngest transposon was used as the overall transposon age (considering it as the point in time where all transposon-components were present).

### Dating of random genomic transposons

For the comparison to human “genomic” transposons (considered were only LINE, SINE, LTR and DNA classes), we merged overlapping transposons from the same species, picked a random sample of 10,000 transposons and applied the same steps as for the enhancers: We prolonged them to 500 bp if they were shorter than that and searched for orthologues in the mentioned vertebrate species and applied phylogenetic reconstruction in the same way as for single-transposon enhancers, as described above.

### Age of enhancer target genes

The target gene age estimates were extracted from the ENSEMBL COMPARA gene phylogenetic trees (ENSEMBL version 94) via the ENSEMBL REST API [66].

### Orthology of target genes and enhancers in human and mouse

We used ENSEMBL Compara to determine the orthologues of human genes in mouse (and vice versa). The pairs of human and mouse genes with an orthology relationship targeted by at least one human and one mouse enhancer constitute the set of common enhancer target genes between the human and mouse correlated enhancer datasets. Enhancer orthologues were determined as liftOver [69] hits.

## SUPPLEMENTARY DATA

Supplementary Data are available online.

