## Supplementary Figures and Tables for "Independent transposon exaptation is a widespread mechanism of shadow enhancer evolution in the mammalian genome"

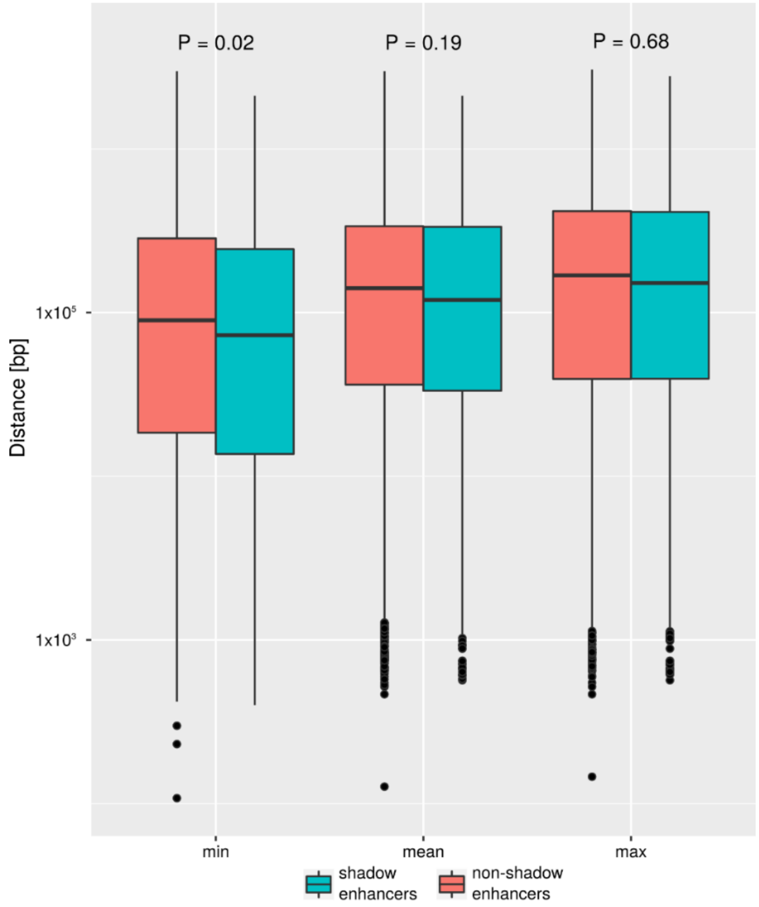


*Supplementary Figure S1. Distance to the (nearest:min, average:mean, farthest:max) TSS of a target gene for 1,280 shadow and 2,243 non-shadow enhancers in the human genome. P-values from Wilcoxon rank-sum tests.*

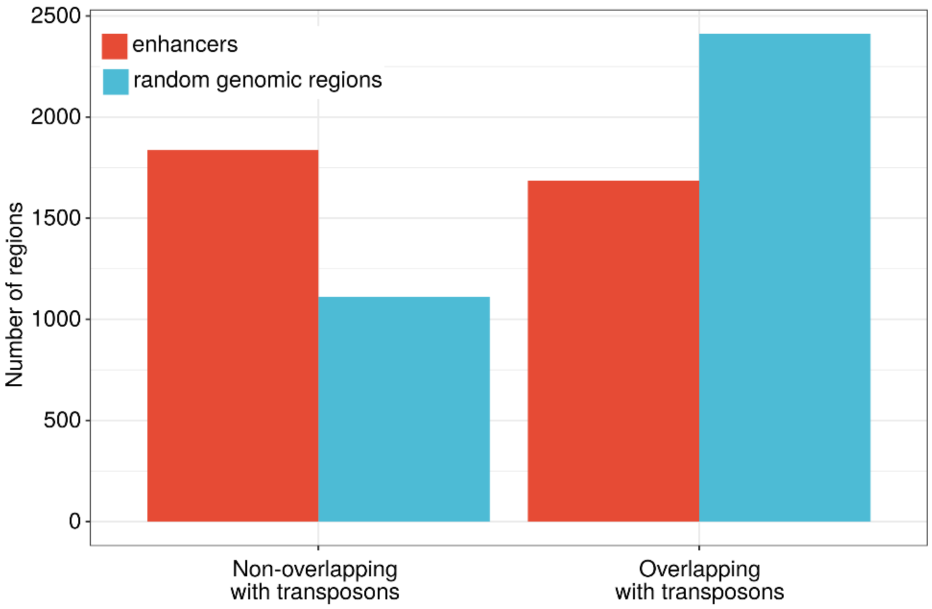


*Supplementary Figure S2. Number of human enhancers annotated as transposons. Random genomic regions show more frequent transposon overlaps than enhancers.*

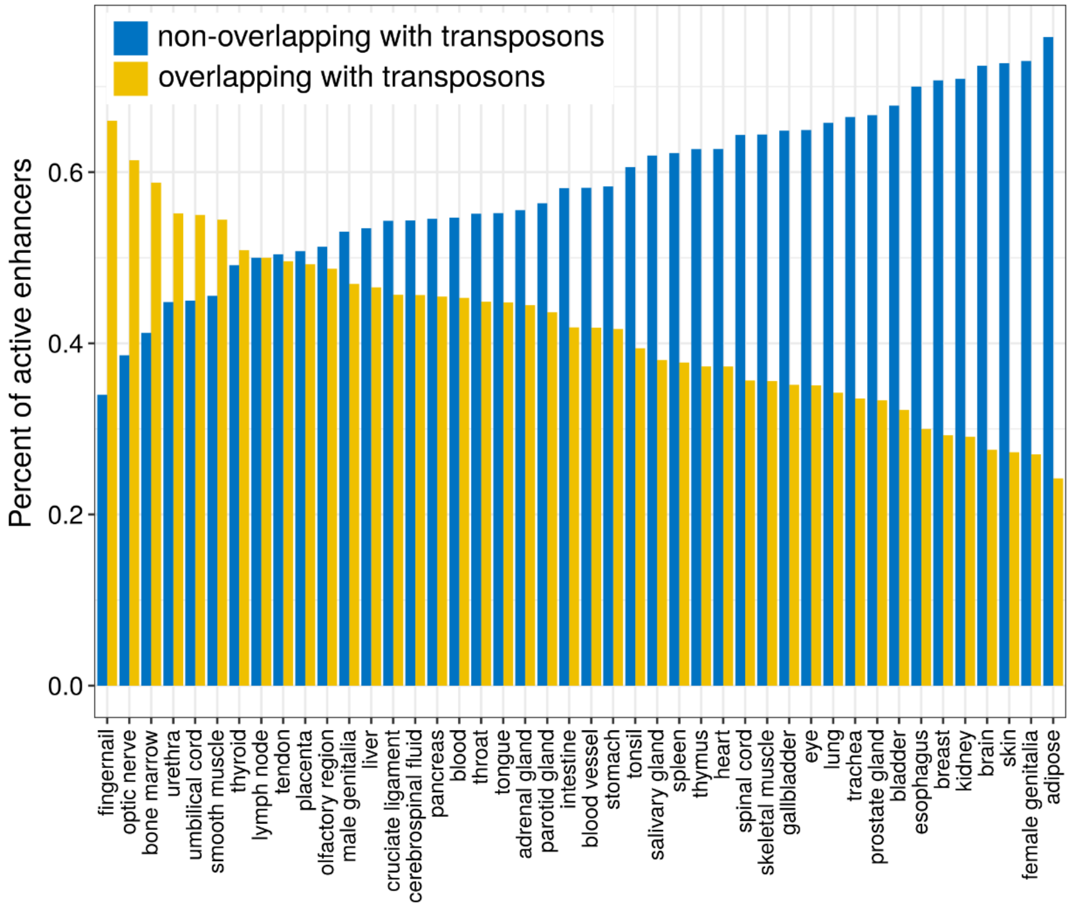


*Supplementary Figure S3. The fraction of transposon overlapping enhancers out of all active enhancers depends on the facet.*

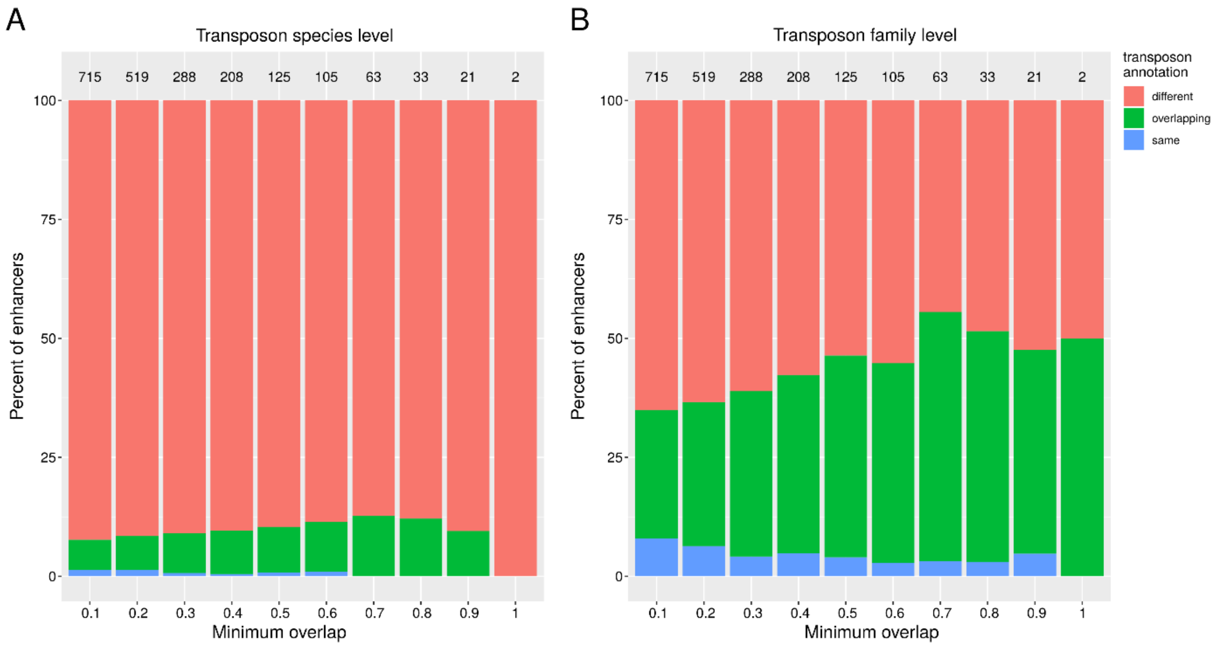


*Supplementary Figure S4. Percent of transposon-shadow enhancer pairs where partners are annotated as transposons from different, shared or identical families, for increasing thresholds of overlap between the enhancer and the transposon sequences for the enhancer to be annotated as a transposon, as fraction of the total enhancer sequence. The numbers above the bars are the total number of pairs where both partners fulfill the overlap criterion.*

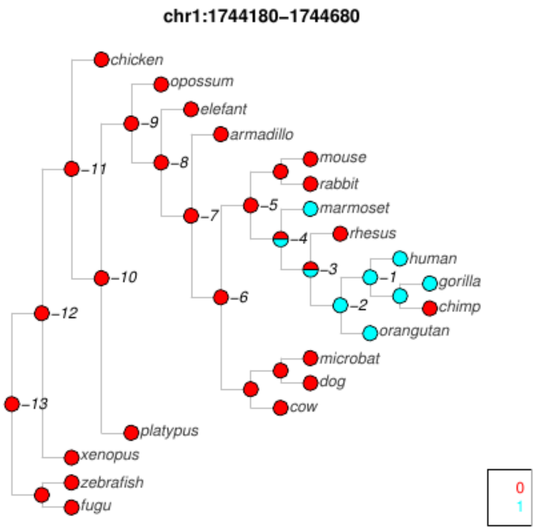


*Supplementary Figure S5. Topology of the phylogenetic tree used for the character state reconstruction of human transposon shadow enhancers. Nodes corresponding to ancestral species are labeled with negative numbers. Ancestral state inferences for the enhancer with coordinates chr1:1,744,180-1,744,680 (hg19). In this example, node -4 is the inferred transposon insertion node.*

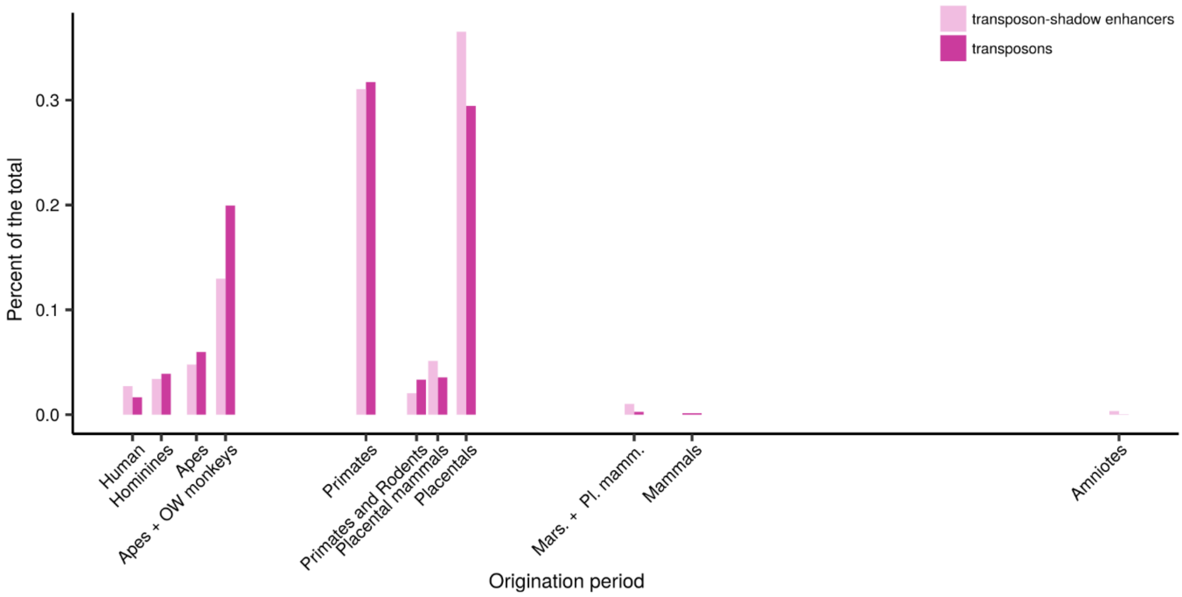


*Supplementary Figure S6. Ages of transposon-shadow enhancers and of random genomic transposon sequences as inferred from the reconstruction of ancestral states in the phylogenetic tree The nodes of the tree were mapped to clades according to the ENSEMBL COMPARA species tree (www.ensembl.org/info/about/speciestree.html).*

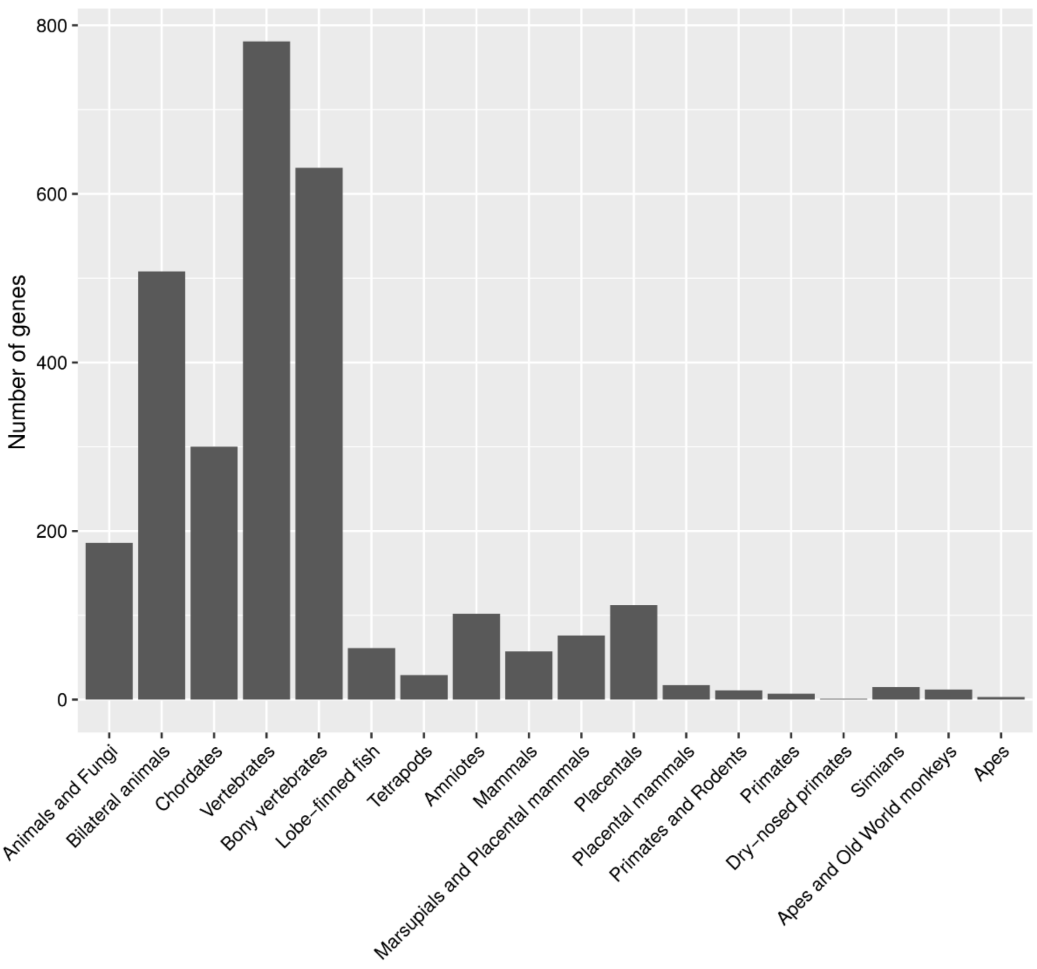


*Supplementary Figure S7. Age of human enhancer target genes. The gene age estimates were extracted from the ENSEMBL COMPARA gene phylogenetic trees. No information was available for 67 genes (excluded from this graph).*

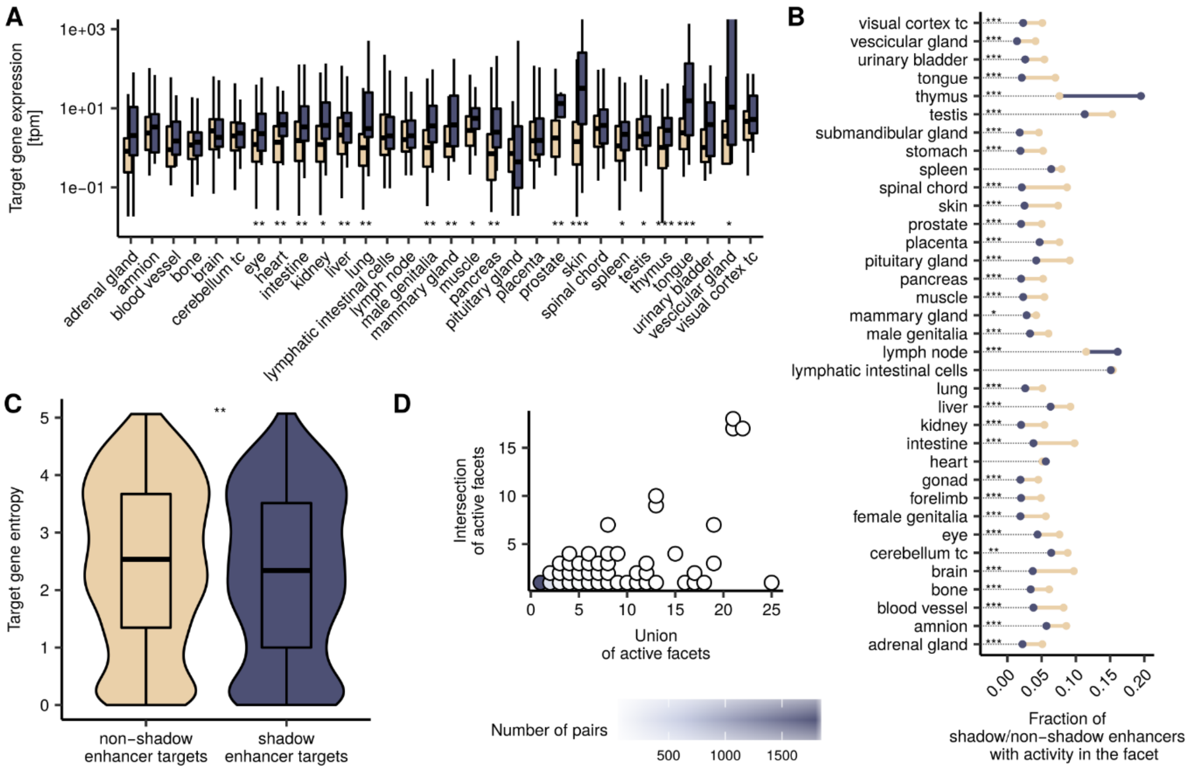


*Supplementary Figure S8. Comparison of shadow and non-shadow mouse enhancers. A) Facet-specific expression of non-shadow (beige) and shadow (purple) enhancer target genes, adjusted P-values of Wilcoxon rank sum tests. '*' P < 0.05, '**' P < 0.01, '***' P < 0.001. B) Dumbbell plot showing the fractions of active non-shadow (beige) and shadow (purple) enhancers per facet. If the fraction of active non-shadow enhancers is larger than that of shadow enhancers, the line between the dots is depicted in beige; otherwise, in purple. The only purpose of the dotted lines is to serve as visual aids. Asterisks indicate adjusted P-values of Fisher's exact tests: Significance code see A). C) Target gene entropies of non-shadow (beige) and shadow (purple) enhancer target genes, Wilcoxon rank sum test. Significance code see A). D) Redundancy of shadow enhancer pairs.*

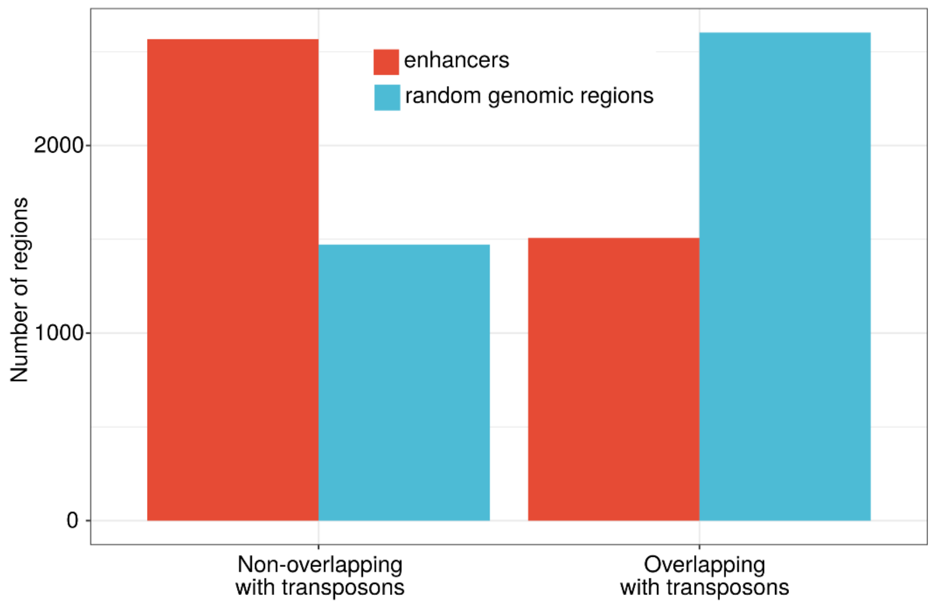


*Supplementary Figure S9. Number of mouse enhancers annotated as transposons. The numbers of elements that overlap with transposons and elements that do not overlap with transposons are reversed in random genomic regions as compared to enhancers.*

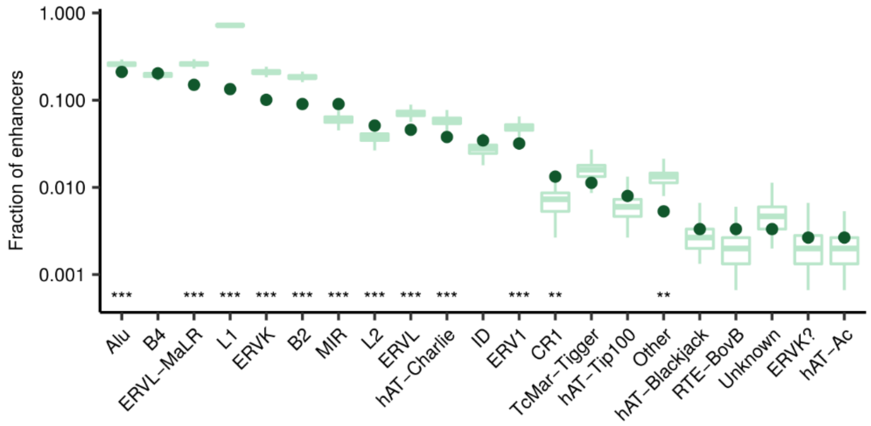


*Supplementary Figure S10. Enrichment of transposon families in transposon enhancers compared to random genomic sequences. Asterisks indicate adjusted empirical P-values. '*' P < 0.05, '**' P < 0.01, '***' P < 0.001. For further description refer to Figure 2A.*

### Supplementary Tables

| | Human | Mouse |
|---|---|---|
| **FANTOM enhancers** | 65,423 | 44,459 |
| - **excluding FANTOM promoters and Ensembl coding exons** | 54,284 | 38,662 |
| - **active in our facets** | 11,582 | 9,426 |
| - **in TADs** | 10,609 | 8,805 |
| - **in TADs that comprise at least one active promoter** | 10,445 | 8,762 |
| - **significantly correlated with one or more promoters** | 3,523 | 4,074 |

*Supplementary Table 1. Construction of the human and mouse enhancer datasets. Number of regions for sets defined based on multiple criteria. Each set of regions is a subset of the set in the previous row.*

| | Human | Mouse |
|---|---|---|
| **FANTOM promoters** | 201,799 | 158,965 |
| - **active in our facets** | 165,749 | 131,995 |
| - **near an Ensembl TSS of a coding gene** | 72,272 | 59,430 |
| - **in TADs** | 60,329 | 52,298 |
| - **in TADs that comprise at least one active enhancer** | 55,612 | 48,323 |
| - **significantly correlated with one or more enhancers** | 6,474 | 8,082 |

*Supplementary Table 2. Construction of the human and mouse promoter datasets. Number of regions for sets defined based on multiple criteria. Each set of regions is a subset of the set in the previous row.*

| **Set** | **Nr of enhancers** | **Nr of pairs** | **Nr of groups** | **Coloring in main figures** |
|---|---|---|---|---|
| **Correlated enhancers** | 4,074 | - | - | Gray |
| **Shadow enhancers** | 1,939 | 2,787 | 670 | Purple |
| **Transposon overlapping enhancers** | 1,507 | - | - | Green |
| **Transposon-shadow enhancers** | 503 | 493 | 192 | Red/magenta |

*Supplementary Table 3. Number of enhancers, enhancer pairs and enhancer groups for multiple sets in mouse. For a description of the sets see Table 1 in the main manuscript..*
